# Investigating Ligand-Mediated Conformational Dynamics of Pre-miR21: A Machine-Learning-Aided Enhanced Sampling Study

**DOI:** 10.1101/2024.06.27.601024

**Authors:** Simone Aureli, Francesco Bellina, Valerio Rizzi, Francesco Luigi Gervasio

## Abstract

MicroRNAs (miRNAs) are short, non-coding RNA strands that regulate the activity of messenger RNAs (mRNAs) by affecting the repression of protein translation, and their dysregulation has been implicated in several pathologies. miR21 in particular has been implicated in tumourigenesis and anticancer drug resistance, making it a critical target for drug design. miR21 biogenesis involves precise biochemical pathways, including the cleavage of its precursor, pre-miR21, by the enzyme Dicer. The present work investigates the conformational dynamics of pre-miR21, focusing on the role of adenine29 in switching between Dicer-binding-prone and inactive states. We also investigated the effect of L50, a cyclic peptide binder of pre-miR21 and a weak inhibitor of its processing. Using time series data and our novel collective variable-based enhanced sampling technique OneOPES, we simulated these conformational changes and assessed the effect of L50 on the conformational plasticity of pre-miR21. Our results provide insight into peptide-induced conformational changes and pave the way for the development of a computational platform for the screening of inhibitors of pre-miR21 processing that considers RNA flexibility, a stepping stone for an effective structure-based drug design, with potentially broad applications in drug discovery.

**TOC Graphic:** 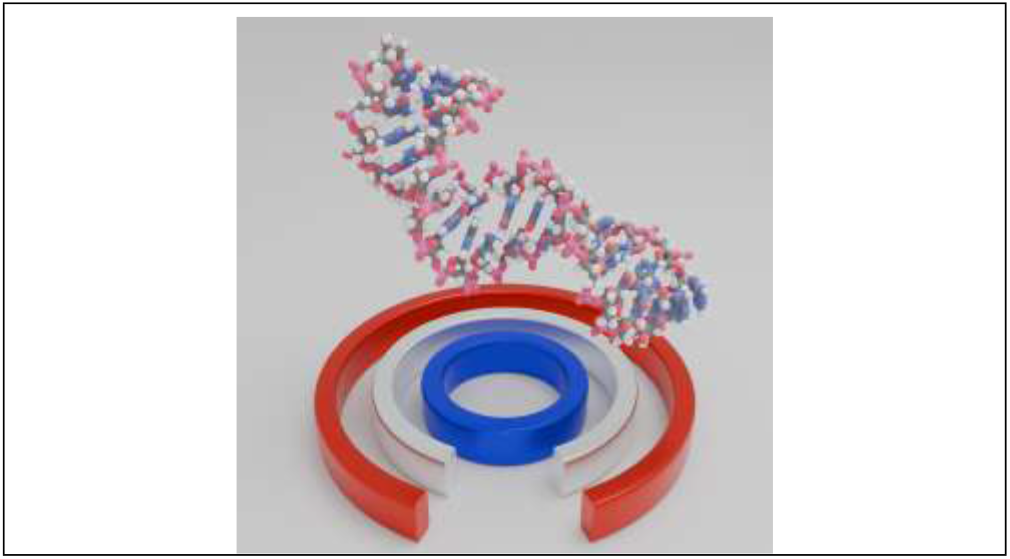

## Introduction

MicroRNAs (miRs) are short, noncoding RNA strands that play a crucial role in posttranscriptionally regulating the messenger RNA (mRNA) activity, leading to mRNA degradation and protein translation repression^1^. MiRs, which typically comprise 18-22 nucleotides, exert significant control over various biological pathways, influencing around 60% of the human genome^2^. Thus, the abnormal dysregulation of miRs is associated with various pathologies, including cancer, metabolic disorders, and cardiovascular diseases^3^. MiRs’s biogenesis follows a precisely tuned biochemical pathway that begins with the miR’ gene transcription into the nucleus (see Fig. 1a)^4^. The RNA polymerase II typically transcribes the miR’s gene into the primary microRNA (pri-miR), a nucleotide RNA stem-loop strand that spans hundreds of nucleotides.

**Figure 1.**
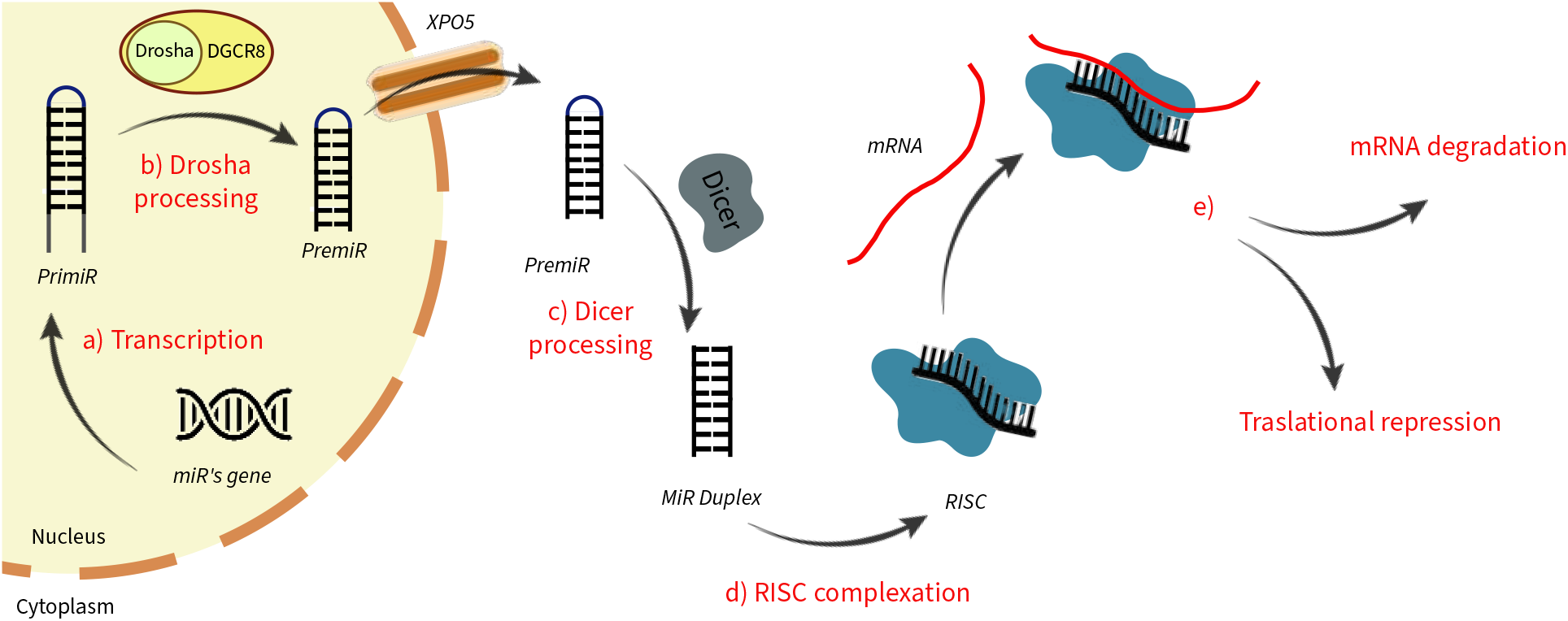
Schematic description of miR’s maturation process. **a)** The pri-miR is generated by RNA polymerase II by transcribing specific miR’s genes on the DNA; **b)** Still in the cell nucleus, the pri-miR interacts with the DCGR8/DROSHA microprocessor that produces the pre-miR; **c)** Once the pre-miR translates into the cytoplasm through the nucleic membrane protein Exportin 5 (EXPO5), it undergoes maturation by interacting with the Dicer enzyme, leading to the fully developed miR duplex; **d)** the miRNA duplex binds one of the AGO protein to product the RISC **e)** the RISC complex selectively recognize the target mRNA leading to its degradation or repressing the translation pathway

Then, the pri-miR is processed by the intranuclear “Drosha/DGCR8” RNase/protein complex. This complex is tasked to recognize and cleave the pri-miR, thus producing a shorter, hairpin-shaped precursor miRNA (pre-miR) of approximately 60-80 nucleotides (see Fig. 1b). The pre-miR is then exported from the nucleus to the cytoplasm by a membrane protein named “Exportin-5” (XPO5). In the cytoplasm, the pre-miR is further processed by the Dicer enzyme (see Fig. 1c) which, in turn, cleaves the pre-miR’s apical loop, and produces a double-stranded RNA duplex, which is composed by one strand known as the “mature miR” and the other one known as the “passenger strand”^5^. The miR duplex is loaded in one of the four “Argonaute” proteins (AGO 1-4) to produce the “RNA-Induced Silencing Complex” (RISC)^6^. Within RISC, the mature miR strand acts as a guide strand that can recognize and bind to specific mRNA molecules possessing a complementary or partially complementary match. The miR-RISC molecular machine either leads to mRNA degradation or represses translation, depending on the degree of complementarity between the miR and the target mRNA.

In this context, microRNA-21 (miR21) emerges as a promising drug target as its abnormal overexpression has been shown to trigger tumorigenesis, sustained cancer growth, and drug resistance across different cell lines^7^. Notably, miR21 exerts an oncogenic effect by downregulating the expression of several tumor suppressor genes (e.g., PDCD4, SPRY2, and PTEN), ultimately fostering uncontrolled cell proliferation and tumor progression^8^. Thus the modulation of miR21 represents a promising therapeutic strategy, with the inhibition of the pre-miR21/Dicer maturation step emerging as one of the most promising and studied approaches^9^. Pre-miR21, a 72-nucleotide stem-loop system, exists in thermodynamic equilibrium between two conformations, i.e. an active state prone to binding and being processed by Dicer, and an inactive one that is naturally

Recent structural investigations have highlighted the crucial role of Adenine29 (A29) in discriminating the two conformations. A29 is a nucleotide close to the Dicer binding site that can adopt either a “stacked-in” or a “bulged-out” orientation leading to the active or the inactive conformation, respectively^13^. Its “bulged-out” orientation may provoke a steric hindrance either preventing Dicer recognition and complexation or destabilizing pre-miR21’s cleavage inside the Dicer machinery^14^. Gaining a detailed understanding of such conformational changes in pre-miR21 would represent the pivotal initial step toward the rational development of effective small molecule inhibitors.

This is why drug discovery campaigns have been in place for the discovery of suitable inhibitors. In recent studies on pre-miR21, docking calculations have been used to accomplish this complex task.^15 16 17^. However, the predictive power of rigid docking methods is rather limited in cases such as pre-miR21 where conformational changes play a relevant role and the changes in the entropy of the system plays a fundamental role. In this regard, the case of L50, a cyclic peptide recently resolved in complex with pre-miR21^18^, is paradigmatic, because it significantly binds to pre-miR21 (*K*_*D*_ *∼* 200*nM*), but it exhibits a poor inhibitory power towards the maturation of pre-miR21 into miR21 (*EC*50 *∼* 10*µM*).

In light of these challenges, we believe that recently developed enhanced sampling and free energy methods might address the challenges posed by the conformational change of pre-miR21, and be used to quantify the effect of L50 on the stabilization of the “bulged-out” configuration. To this end, we initially simulated the “stacked-in/bulged-out” equilibrium of A29, followed by MD simulations of the L50/pre-miR21 complex. To estimate the free energy of the conformational change, the timescale accessible by standard MD simulations is not sufficient, therefore we employed Collective Variable (CV) based enhanced sampling techniques^19,20^. Specifically, we used OneOPES, a novel replica-exchange sampling scheme designed to ease the exploration of hidden yet relevant degrees of freedom^21^. OneOPES leverages a temperature gradient along the replicas’ ladder and employs several different CVs simultaneously, allowing for an accurate and efficient estimate of free-energy differences. The proposed approach proved to be able to discern the subtle conformational changes and their associated energetics between the apo pre-miR21 and the L50/pre-miR21 complex, providing valuable insights about how to induce structural alterations of pre-miR21 and to modulate its physiological activity.

Our study provides a robust computational platform for investigating conformational changes in miRs also in the presence of a ligand and estimating the corresponding free-energy change. We propose that our protocol serves as a tool for screening and prioritizing potential inhibitors of pre-miR21 processing, offering a cost-effective and time-efficient alternative to traditional experimental approaches. Furthermore, our procedure can be considered a blueprint for studying ligand-induced conformational changes in other protein or nucleic acid targets. By measuring the energetics associated with the activation of the apo state and comparing it to the free-energy surface obtained in the presence of a ligand (either a peptide or a small molecule), our methodology provides a comprehensive understanding of ligand-binding events and their impact on target dynamics. Overall, our computational framework holds promise for accelerating drug discovery efforts and advancing our understanding of molecular interactions in complex biological systems.

## Methods

### MD simulations

The apo- and holo-structures of pre-miR21 were retrieved from PDB ID: 5UZT and 5UZZ^18^, respectively. In particular, the apo structure displays the apical loop of pre-miR21 in its “stacked-in” conformation whereas the holo structure (in the “stacked-in” conformation as well) is found in complex with L50 (i.e., cyclo(RVRTRGKRRIRRpP)), a cyclic peptide spanning 14 residues and characterized by a d-Proline in position 13. The structures thereby obtained were embedded into a tailored octahedral box, solvated with TIP4PW water model, and neutralised with Na^+^ ions. A second replica of the holo system was built with a salinity of 0.15 NaCl, to investigate the effect of different ionic strengths on L50’s binding stability. The reparametrised Amber14ff force field was employed^22^ in the MD engine GROMACS 2023^23^. Each simulation box underwent a thermalisation cycle using restraints on heavy atoms with the following protocol: 100 ns of NVT simulation followed by 100 ns of NPT simulation for each temperature at 300 K. The particle-mesh-Ewald (PME) method was used to treat the electrostatic interaction^24^. On the van der Waals interactions, a cut-off distance of 1.0 nm was applied. The pressure was fixed at a reference value equal to 1 bar thanks to the C-rescale barostat^25^, whereas the temperature was controlled through the Nose-Hoover thermostat^26^.

### OneOPES MD simulations

The conformational change of apo- and holo-pre-miR21 was investigated through CV-based enhanced sampling methods. Notably, we resorted to employing the OneOPES sampling scheme^21^, a derivative technique of the “On-the-fly probability enhanced sampling” (OPES) algorithm in its “Explore” flavor^27^. In OneOPES, a replica-exchange framework of 8 independent trajectories is set up to ensure the exploration and the convergence of the FES under investigation. Such replicas are divided into two groups, i.e., a convergence-dedicated replica (replica 0) and seven exploratory trajectories (replicas 1-7). They all share OPES Explore as the main sampling engine carried out on a set of leading CVs. Replicas 1-7 are progressively heated (up to 320 K) thanks to OPES Expanded (OPES MultiT, hereafter), to ease overcoming hidden degrees of freedom^28^. In the present scenario, we resorted to tailored CVs to drive pre-miR21’s conformational transition. In detail, we employed the “Harmonic Linear Discriminant Analysis” (HLDA) CV, a weighted linear combination of 10 intraRNA contacts able to maximise the pairwise discriminatory power among the two pre-miR21’s states^29^. The contacts and the corresponding coefficients are shown in Tab. 1:

**Table 1:**
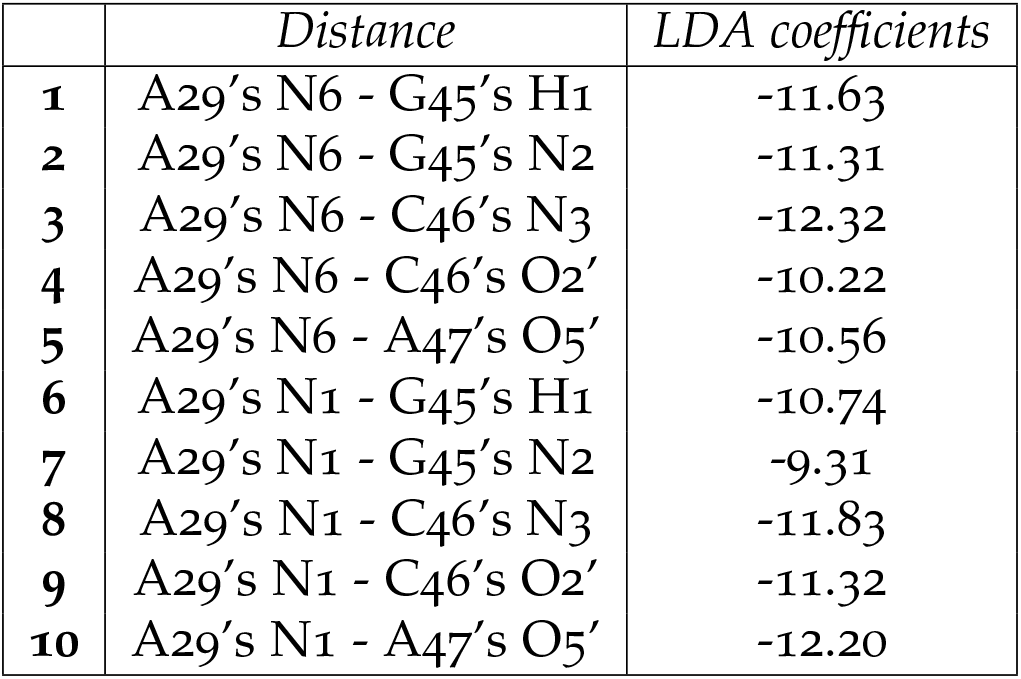
Table reporting the 10 intraRNA contacts and the corresponding LDA coefficients.

|  | *Distance* | *LDA coefficients* |
| --- | --- | --- |
| **1** | A29's N6 - G45's H1 | -11.63 |
| **2** | A29's N6 - G45's N2 | -11.31 |
| **3** | A29's N6 - C46's N3 | -12.32 |
| **4** | A29's N6 - C46's O2' | -10.22 |
| **5** | A29's N6 - A47's O5' | -10.56 |
| **6** | A29's N1 - G45's H1 | -10.74 |
| **7** | A29's N1 - G45's N2 | -9.31 |
| **8** | A29's N1 - C46's N3 | -11.83 |
| **9** | A29's N1 - C46's O2' | -11.32 |
| **10** | A29's N1 - A47's O5' | -12.20 |

**Table 2:** Table depicting the different CVs and parameters used for the OneOPES simulations. On the rows “OPES Explore” and “OPES MultiCV” we report how the CVs have been arranged along the replicas. On the row “OPES MultiT”, we display the highest temperature achieved along the trajectory starting from the temperature of the thermostat (i.e., 300 K).

| Replicas | 0 | 1 | 2 | 3 | 4 | 5 | 6 | 7 |
| --- | --- | --- | --- | --- | --- | --- | --- | --- |
| OPES Explore | HLDA | HLDA | HLDA | HLDA | HLDA | HLDA | HLDA | HLDA |
| OPES MultiCV 1 | - | w0 | w0 | w0 | w0 | w0 | w0 | w0 |
| OPES MultiCV 2 | - | - | t2 | t2 | t2 | t2 | t2 | t2 |
| OPES MultiCV 3 | - | - | - | t3 | t3 | t3 | t3 | t3 |
| OPES MultiCV 4 | - | - | - | - | t4 | t4 | t4 | t4 |
| OPES MultiCV 5 | - | - | - | - | - | t5 | t5 | t5 |
| OPES MultiCV 6 | - | - | - | - | - | - | t6 | t6 |
| OPES MultiCV 7 | - | - | - | - | - | - | - | t7 |
| OPES MultiT | - | 302K | 304K | 306K | 308K | 310K | 314K | 320K |

For a complete description of the atoms involved in HLDA, please refer to Fig. S4a.

#### Auxiliary CVs have been introduced on exploratory trajectories, to increase their sampling power

- **w0:** water coordination site around A29 (i.e. the “Hydration” CV)^30 31^;
- **t2-7:** nucleotides’ torsional angles.

In this framework, the frequency of exchange among the replicas was 2000 integration steps. Replicas 0-7 contain a layer of OPES Explore with a bias of 20.0 kJ/mol and a PACE (i.e., frequency of bias deposition) of 20000 steps. Moreover, replicas 1-7 underwent additional OPES Explore layers (OPES MultiCV) with a bias of 3.0 kJ/mol on the auxiliary CVs and a PACE of 40000 steps.

To run the simulations the MD engine GROMACS 2023 patched with PLUMED 2.9 was employed^32^. Regarding the thermostat and the barostat, we used the same protocol described in the section “MD simulations”.

### Cluster analysis

Cluster analyses on the MD trajectories were performed using GROMACS’s *gmx cluster* routine, using the *gromos* algorithm. The cluster families of pre-miR21 in the apo- and holo-pre-miR21 MD simulations were obtained by aligning the trajectory on the P, C4’, and C5’ atoms of pre-miR21’s paired nucleotides, and computing the RMSD among the same sample of atoms. An RMSD threshold value of 2.0 Å was selected considering the number of generated cluster families and the similarity of RNA conformations within a cluster family.

### Binding interface evaluation

To assess the interactions in the L50/pre-miR21 complex, we utilised the PLOT NA routine of the “Drug Discovery Tool” (DDT) to estimate the frequency of occurrence of contacts^33^. Similar to previous studies, we defined a neighbouring cutoff value of 4.0 Å between two interacting residues to analyse the binding interfaces.

### Cross-correlation analysis

To evaluate the correlated motions between nucleobases in the apo- and holo-pre-miR21 systems, we utilized cross-correlation analysis (also known as Pearson-correlation coefficient analysis)^34^. An in-house Python script was employed to calculate the correlation matrix for each pair of residues using the following formula:

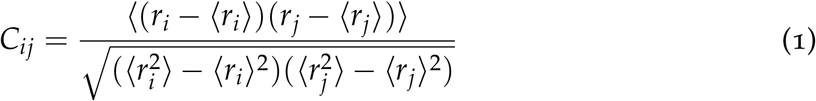

where *r*_*i*_ and *r*_*j*_ are the position vectors of the *Cα* atoms in residues *i* and *j*, respectively. The angle brackets denote time averages computed along the simulations. The final *C*_*ij*_ value ranges from -1.0 to 1.0.

## Results

Our investigation focused on elucidating the conformational dynamics underlying the activation of pre-miR21, an RNA molecule that is crucial in regulatory processes. Our primary aims were to characterize the structural transition associated with pre-miR21 activation and map its corresponding free-energy landscape, both in its apo form and in the presence of the cyclic peptide L50. To achieve these goals, we employed a comprehensive approach combining classical MD simulations with state-of-the-art data-driven enhanced sampling techniques. Specifically, we conducted atomistic MD simulations spanning a total cumulative time of 1200 ns on the pre-miR21 molecule. Concurrently, we investigated the binding complex L50/pre-miR21 (*holo-pre-miR21* hereafter) through an analogous simulation protocol. The following sections present a detailed exposition of our findings, shedding light on the conformational dynamics of pre-miR21 activation.

### Structural properties of pre-miR21

The initial structure of both apo- and holo-pre-miR21 (Fig. 2a and Fig. 2b, respectively) underwent 800 ns of classical MD calculation to explore the conformational plasticity of the RNA strand and the impact of the L50 peptide. Root Mean Square Deviation (RMSD) analysis of the backbone atoms of the paired nucleobases was conducted to inspect the nucleotide’s overall dynamics. As depicted in Fig. 2c, apo-pre-miR21 exhibited rather high average RMSD values (*∼*2.0 Å), suggesting a poor degree of conformational stability. This observation was further corroborated by cluster analysis, revealing a significant number of distinct conformational states explored by the nucleotides during MD simulations (see “Methods” section for further details). However, a few key conformational differences emerge when a deeper analysis of the nucleobases is performed. Indeed, the two most populated conformation families present a different position of A29, with the largest cluster characterized by A29 in the “bulged-out” state, and the second largest cluster presenting A29 in the “stacked-in” conformation (see Fig. S1a). This observation indicates that A29 and its surrounding region have a high level of mobility, which might be functional to adapt pre-miR21’s shape to the binding partner Dicer during the processing of the miR duplex. Conversely, a different scenario emerges when performing the same analysis on holo-pre-miR21. The average RMSD value remained stable at *∼*1.0 Å throughout the MD simulation (see Fig. 2d), indicating a significant reduction in the flexibility of the RNA strand. A similar picture arises from the cluster analysis, which revealed the presence of only a single largely populated cluster over the whole trajectory, characterized by A29 in the “stacked-in” state (see Fig. S1b). Additionally, we extended the RMSD analysis to the apical loop of pre-miR21, showing that the inhibition exerted by L50 on pre-miR21’s conformational flexibility propagates to the apical loop as well. As displayed in Fig. S2a and S2b, the apical loop in the apo-MD simulation exhibited a wide range of RMSD values throughout the whole trajectory, with fluctuations reaching up to 12.0 Å. Conversely, the same apical loop displayed significantly lower fluctuations in holo-pre-miR21, stabilizing at an RMSD value of *∼*4.0 Å. This further supports the observation that L50 restricts the overall conformational freedom of pre-miR21, not only around A29 but also in the distant apical loop region. For additional analyses on pre-miR21’s and L50’s behaviour, please see Fig. S2.

**Figure 2.**
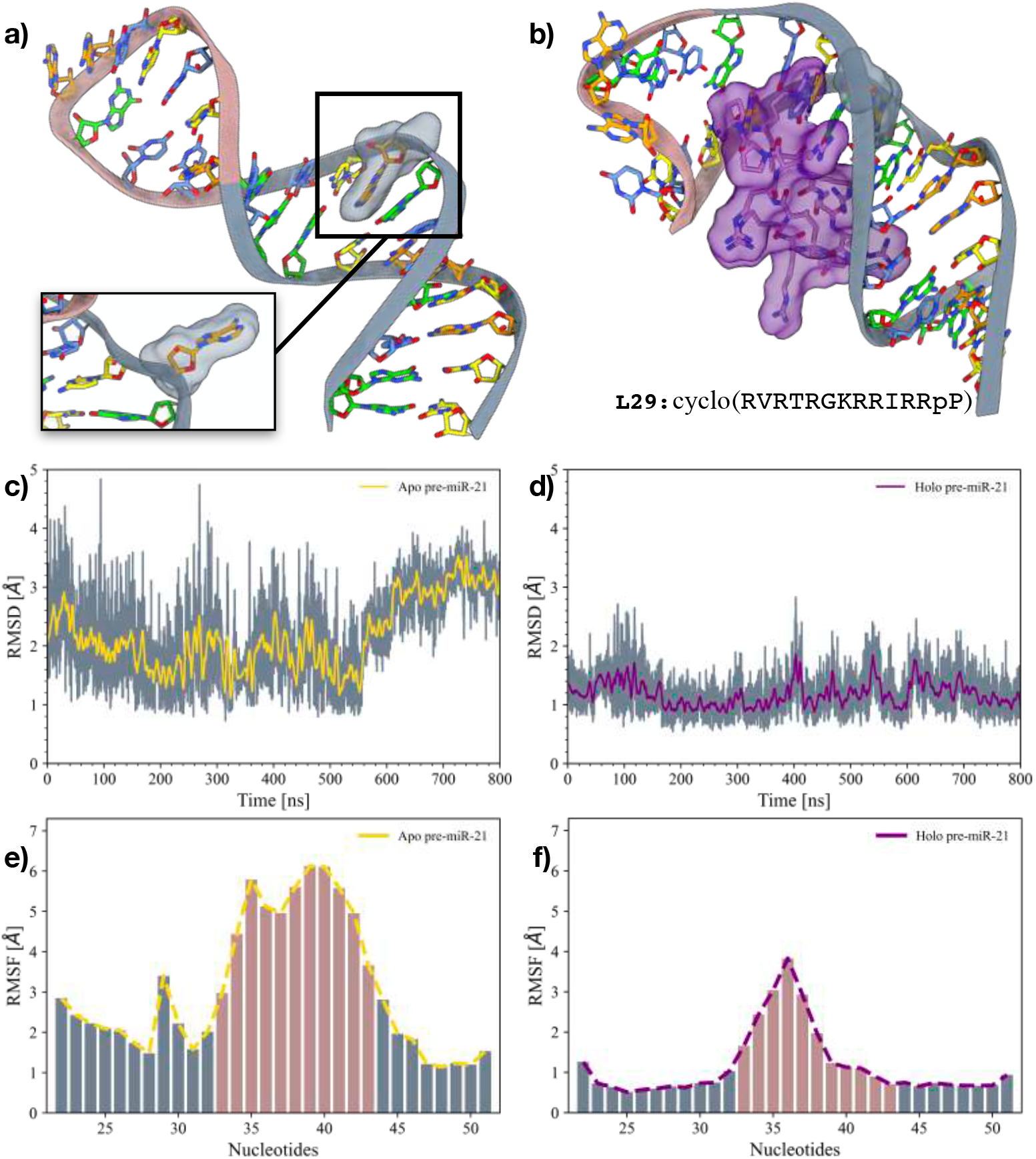
Computational investigations on the apo- and holo-pre-miR21 systems. **a)** 3D structure of apo-pre-miR21. The inset displays a snapshot from a frame when A29 assumes the “bulged-out” state. Adenines are colored in orange, guanines in green, cytosines in yellow, and uracils in azure. The phosphate backbone of the nucleotides assuming a proper Watson-Crick pairing is colored in grey, whereas it is colored in pink for uncoupled nucleobases. A29 is highlighted through its solvent-exposed surface. **b)** 3D structure of L50 bound to pre-miR21. Nucleobases are colored as in panel (a). L50 and A29 are highlighted through their solvent-exposed surface. The primary sequence of L50 is reported at the bottom of the panel. **c-d)** RMSD of pre-miR21’s backbone in the apo- (c) and holo- (d) 800 ns MD simulations. The RMSD running averages are superimposed in yellow and purple, respectively. **e-f)** Histograms displaying the per-nucleobase RMSF for the apo- (e) and holo-pre-miR21 (f) systems. The nucleotides have been colored as in panel (a).

A further indication of the high flexibility of pre-miR21 emerges from the evaluation of the Root Mean Square Fluctuation (RMSF). Such analysis indicated that the region surrounding A29 in apo-pre-miR21 exhibits higher plasticity with respect to other paired nucleobases, supporting our hypothesis regarding localized conformational dynamics. Additionally, a comparison of RMSF values between the apo- and holo-simulations provides further evidence of the suppression of pre-miR21’s conformational plasticity. In the apo-conformation, the entire RNA strand demonstrated the highest flexibility, with a notable peak in RMSF centered around A29, measuring *∼*3.5 Å. This flexibility is markedly reduced in the holo-conformation, where the RMSF value around A29 decreases to *∼*0.5 Å. While this investigation allowed us to identify specific differences between the apo- and holo-systems, more in-depth analyses were necessary to properly assess the overall stabilization induced by the presence of L50.

### Effect of L50 binding on pre-miR21’s dynamics

To elucidate the effect of peptide binding on the functional dynamics of pre-miR21, we performed a cross-correlation analysis (CCA) on the nucleobases in both the apo- and holo-forms. Indeed, such a method allows for identifying short- and long-range allosteric effects between different parts of the RNA strands, putting into evidence the internal communication networks^35^. Namely, we focused our analysis on 4 macro-regions of pre-miR21 defined as follows:

- *“G22-U26”*: the 5’ end of pre-miR21 (red in Fig. 3a);
- *“G28-U31”*: where A29 is located (yellow in Fig. 3a);
- *“U33-A36”*: the unstructured loop of pre-miR21 (blue in Fig. 3a);
- *“C39-A50”*: a wide portion of the RNA hairpin complementary to the regions mentioned above (green in Fig. 3a).

**Figure 3.**
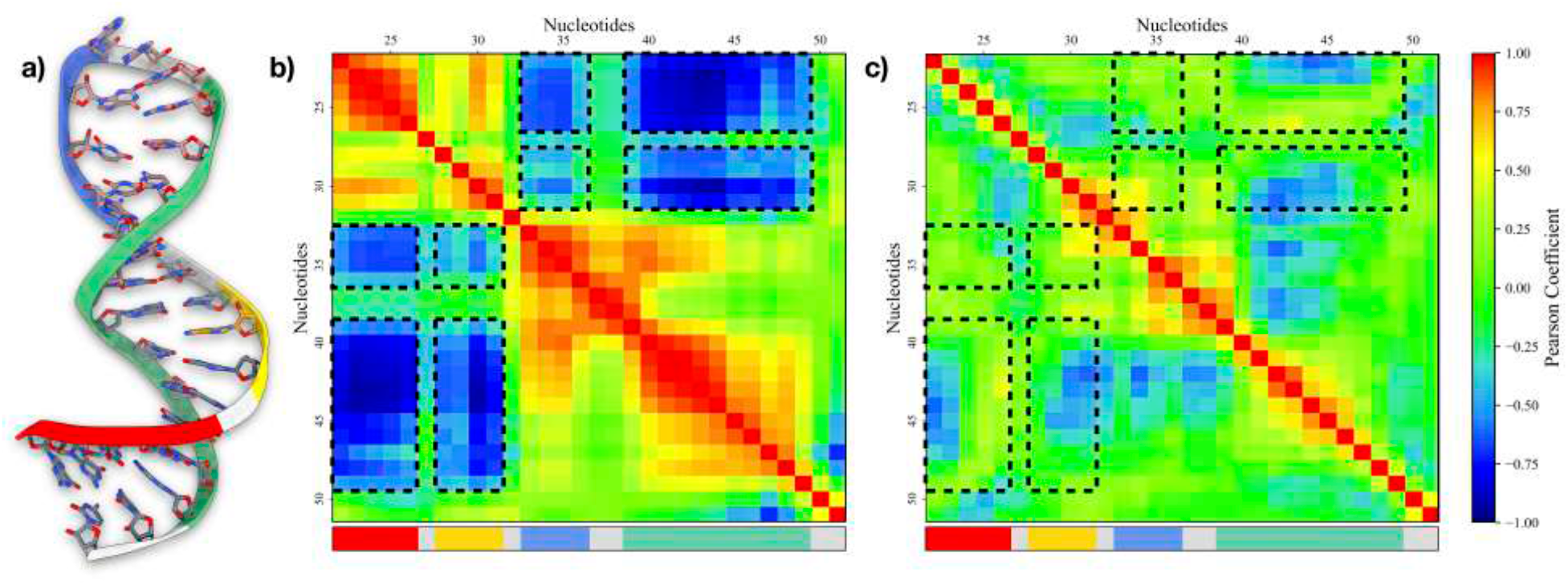
Studies on the conformational flexibility of pre-miR21. **a)** Schematic representation of the major regions composing pre-miR21. **b-c)** Pearson coefficients computed between pairs of residues in apo-pre-miR21 (b) and holo-pre-miR21 (c), respectively. The two maps are colored according to the color bar on the right side. Areas displaying high linear correlation are indicated as black dashed line squares. Additionally, at the bottom of both maps, a rectangular box has been added, colored according to the macro regions identified on pre-miR21.

Regarding apo-pre-miR21, regions “G22-U26” and “G28-U31” show anti-correlated motions (e.g. significant negative Pearson coefficients) with both the regions “U33-A36” and “C39-A50” (see Fig. 3b). This suggests a coordinated physiological movement among the different areas of pre-miR21, which may be essential for its binding to the Dicer protein and subsequent maturation into the miR duplex. Upon binding with the cyclic peptide L50, the internal interaction network of pre-miR21 is significantly suppressed, leading to a notable reduction in its conformational flexibility (see Fig. 3c). Indeed, region “G22-U26” shows no correlation at all with the other pre-miR21’s areas, whereas “G28-U31” displays anti-correlation only with “C39-A50”, even though of only limited intensity. To better understand the inhibitory effect of L50 on pre-miR21, we also performed a principal component analysis (PCA) on the motion of the RNA strand. Projecting the first eigenvector for the apo-pre-miR21 system, it becomes evident that the main motion involves regions “G28-U31” and “C39-A50”, which move in opposite directions (see Fig. S3a). In contrast, L50 binding causes a completely different motion in holo-pre-miR21, where the first eigenvector is dominated by the movement of the macro-region “U33-A36” (see Fig. S3b). This suppression of pre-miR21’s conformational plasticity may have important implications for the molecule’s stability and its subsequent interactions, potentially affecting its role in the maturation process and its binding affinity with other proteins such as Dicer. Even though classical MD simulations provided us with some insights into the mechanism of action of L50, achieving a complete description of the conformational changes in pre-miR21 proved challenging due to timescale limitations. Consequently, we decided to proceed with enhanced sampling simulations on both the apo- and holo-systems to gain a more comprehensive understanding.

### OneOPES simulations

While our previous insights showed promise, they were still constrained by the timescale problem, as it often happens in MD simulations. For instance, it was not possible to observe any “stacked-in/bulged-out” conformational changes in the holo-pre-miR21 simulations, an event that would physiologically happen but typically on a timescale inaccessible by plain MD. To achieve a comprehensive description of the conformational landscape available to pre-miR21, we employed OneOPES, a novel replica-exchange sampling scheme deputed to ease the overcoming of hidden energy barriers and, thus, deliver the full reconstruction of the free-energy surface^21,36^. Developed as a combination of a number of variants of the “On-the-fly Probability Enhanced Sampling” (OPES) technique^27^, it relies on a set of reaction coordinates named “Collective variables” (CVs) that characterize the process under investigation and a bias potential that guides the system along the desired pathways.

In the present scenario, two 400ns-long OneOPES calculations were carried out using 8 different replicas. As CVs, we resorted to employing a “harmonic linear discriminant analysis” (HLDA) CV, built as the linear combination of 10 intranucleobase contacts^29^ (see “Methods” and Fig. S4a for additional details). This HLDA CV was able to track the conformational change associated with pre-miR21’s A29 (see Fig. S4b and S4c), revealing a significant alteration in the FES between apo- and holo-pre-miR21. As displayed in Fig. 4a, the apo-pre-miR21’s FES shows 2 energy minima located at HLDA*∼*-30 and HLDA*∼*-90. The former is the lowest free energy basin, corresponding to the “stacked-in” state. In contrast, the latter is the state with the highest energy value, representing the “bulged-out” state. We measured the free energy difference of the minima over time and, after *∼*100ns of simulation, it converged to be *∼*-2.6 kcal/mol (see Fig. S2b-c).

**Figure 4.**
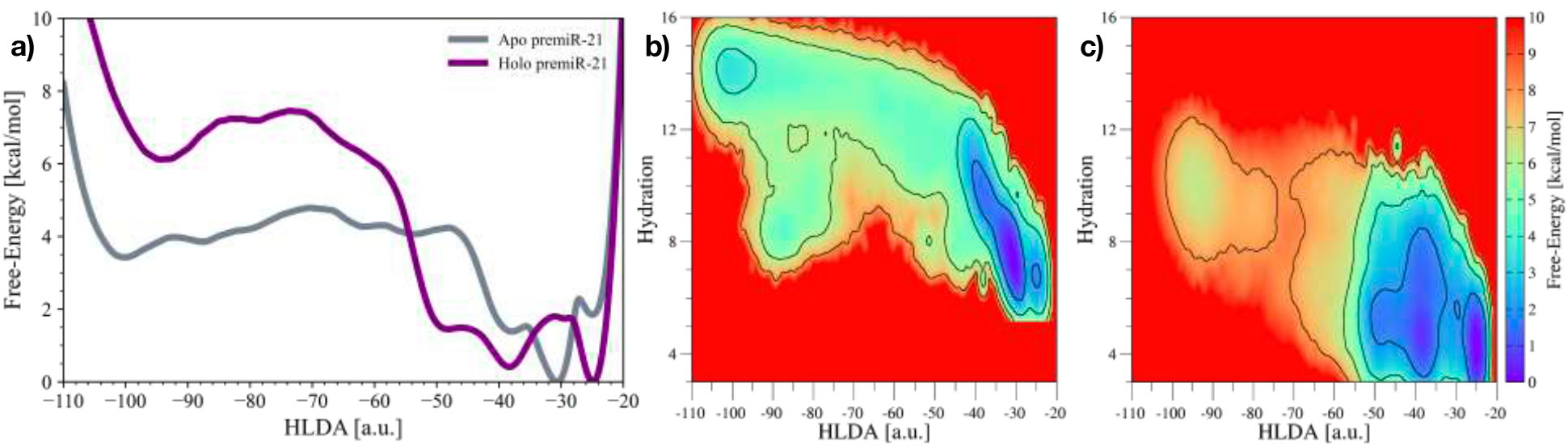
Comparison of OneOPES simulations on the apo-pre-miR21 and holo-pre-miR21 systems. **a)** 1D free-energy profiles associated with apo-pre-miR21’s and holo-pre-miR21’s “stacked-in/bulged-out” conformational change. **b-c)** 2D free-energy surfaces depicting the conformational transition in the apo-(b) and holo-pre-miR21 (c) systems. For both OneOPES simulations, the accumulated bias potential has been reweighted on both HLDA and Hydration CVs.

A rather different scenario was obtained for the holo-complex. Indeed, the presence of the L50 peptide stabilises the free-energy basin corresponding to the “stacked-in” state, slightly disfavouring the Dicer-prone “bulged-out” conformation. After *∼*200ns of simulation, the difference in free-energy was estimated to be *∼*-5.6 kcal/mol (see Fig. S3a-b). A better understanding of the mechanism governing this conformational change was achieved by re-weighting the collected bias potential on both HLDA and “Hydration shell” CVs (see Fig. 4b and Fig. 4c). The latter CV monitors the amount of water surrounding A29 (see details in “Methods” section). Regarding the apo-system, the 2D FES clearly shows that the transition between the “stacked-in” and “bulged-out” states occurs with a significant increase in the hydration level of this nucleobase. In particular, two different basins have been identified for the “bulged-out” state, i.e. the fully-hydrated canonical one (Hydration *∼*15) and an auxiliary one (Hydration *∼*9) only partially hydrated.

Conversely, the holo-pre-miR21 system is characterised by an overall less hydrated conformation pool, mainly due to the presence of the large cyclopeptide L50 bound to the RNA hairpin. While hydration has a minimal effect on the “stacked-in” state, the “bulged-out” one is way more influenced. Indeed, only the partially hydrated “bulged-out” state has been measured to be a minimum in the FES, whereas the canonical fully hydrated conformation is a high energy state. These findings may elucidate why L50 is a good binder to pre-miR21, but a very weak inhibitor (*EC*_50_ = 10*µM*)^18^. Indeed, while L50 is capable of disrupting the physiological conformational plasticity of pre-miR21 and impacting downstream miR21 biogenesis, it fails to shift the “stacked-in/bulged-out” equilibrium in favor of the “bulged-out” state, thereby limiting its inhibitory effec-tiveness.

### The bulged-out state of pre-miR21 in presence of L50

The impact of L50 on the pre-miR21 conformational dynamics underscores the intricate regulatory mechanisms governing miRNA activation. By modulating the plasticity of the overall RNA strand, ligands such as L50 possess the capacity to down-regulate Dicer binding and consequently inhibit miR21 processing.

Nevertheless, its discovery was due to the systematic screening of a large peptide library originally designed to bind *“Bovine Immunodeficiency Virus Trans-Activation Response element”* (BIV TAR), without carrying out a dedicated structure-based drug design campaign due to the lack of a pre-resolved structure^18^. Thanks to our holo-pre-miR21 OneOPES simulations, we could extract the portion of the trajectory corresponding to the “bulged-out” state, which has not been experimentally resolved so far. As displayed in Fig. 5a, a cluster analysis carried out on such a conformation pool revealed the presence of a single largely populated cluster, whose centroid is shown in Fig. 5b. In this regard, it is worth mentioning the contacts established by L50’s Arg1 and pre-miR-21, where the amino acid engages a *π −π* stacking with A29 and, at the same time, builds a salt bridge with G28’s phosphate group. A full list of the L50/pre-miR21 interactions is reported in Tab. S1, whereas a full comparison with the “bulged-out” conformation extrapolated from apo-pre-miR21 OneOPES simulation is reported in Fig. S6. The structure displayed in Fig. 5b is released as a PDB in the supplementary material. It can be used as a starting point to develop novel small molecules or peptides capable of both disrupting pre-miR-21’s conformational plasticity and shifting A29’s conformational equilibrium towards the “bulged-out state”. In our opinion, this approach has the potential to lead to the development of potent inhibitors of miR-21 processing.

**Figure 5.**
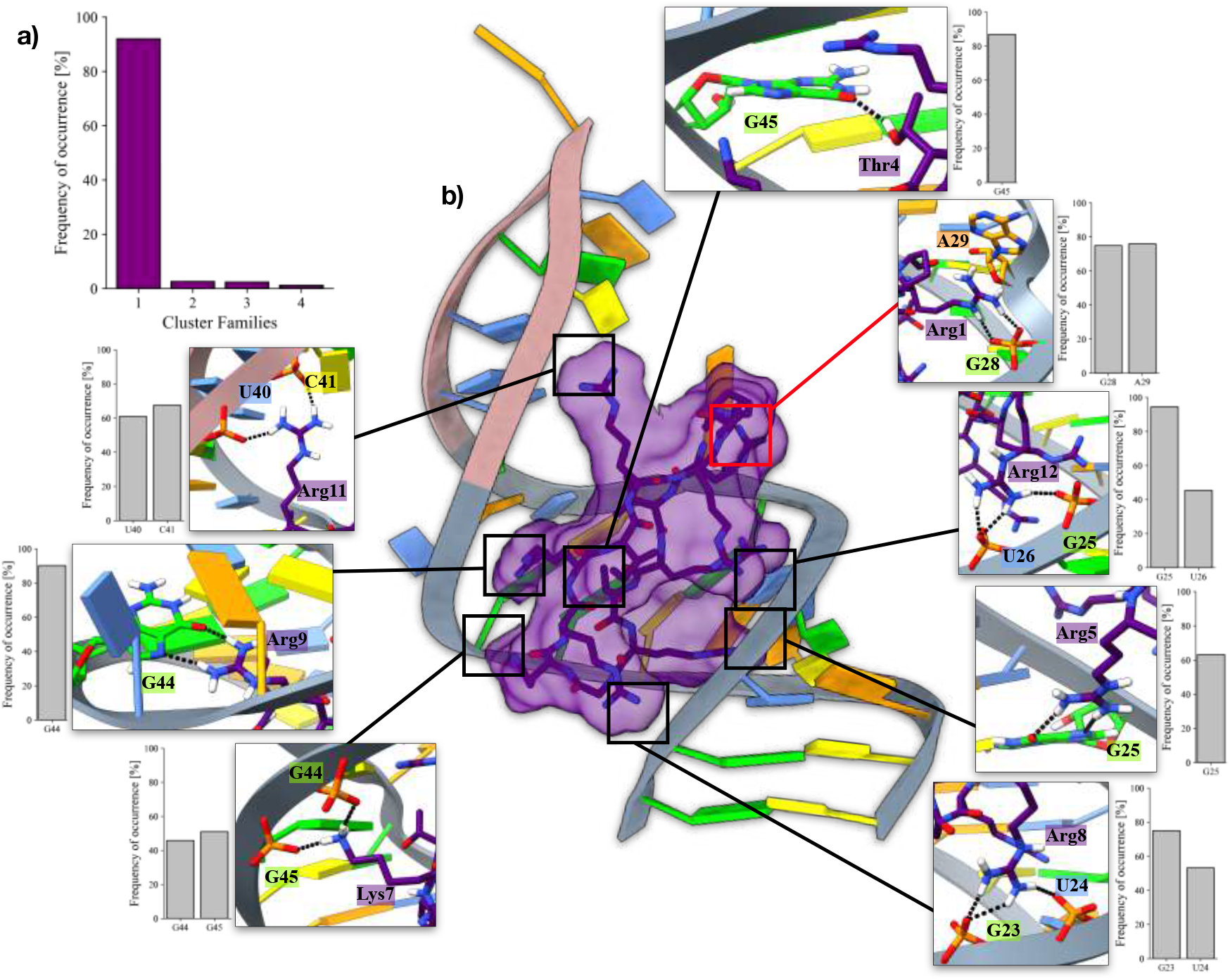
Bulged-out state extrapolated from the holo-pre-miR21’s OneOPES simulation. **a)** Histogram reporting the results of the cluster analysis carried out on the “bulged-out” conformation pool generated through the OneOPES simulation on the holo-pre-miR21 system. **b)** Structure of the centroid associated with the most populated cluster family. The insets display the interactions established by L50’s residues with pre-miR21 and their frequency of occurrence. The red box highlights the interactions established by A29. Adenines are colored in orange, guanines in green, cytosines in yellow, and uracils in azure. The phosphate backbone of the nucleotides assuming a proper Watson-Crick pairing is colored in grey, whereas it is colored in pink for uncoupled nucleobases. L50 is colored in purple and highlighted through its solvent-exposed surface.

## Conclusions

MiRs play pivotal roles in regulating gene expression and influencing diverse cellular processes, making them attractive targets for therapeutic intervention^37^ and subject to an intense recent research effort^38–40^. Dysregulation of miR function has been implicated in various diseases, highlighting the potential of miR-based therapeutics in precision medicine. However, significant changes in the conformation of the miRNA pose substantial challenges to classical rigid docking approaches for computer-aided drug discovery. In the present study, we employed a novel enhanced sampling protocol named OneOPES to investigate the conformational dynamics of pre-miR21 and the impact of ligand binding on its activation process. By exploiting the unique features of OneOPES (e.g. multiCVs, thermal ramp, replica exchange), we could uncover insights into the conformational change governing pre-miR21 activation, highlighting the intricate interplay between structural dynamics and functional outcomes.

Our simulations confirm the inherent conformational plasticity of pre-miR21, revealing a high degree of flexibility in its backbone structure while highlighting localized dynamics within key nucleotide residues. Notably, the conformational equilibrium between the “stacked-in” and “bulged-out” states of A29 emerged as a critical aspect of pre-miR21’s physiological activity, assuming a relevant role in the context of Dicer binding and miR processing. In parallel, we have provided mechanistic insights into the ligand-induced modulation of pre-miR21 conformational dynamics. In this regard, we studied the effect of the cyclic peptide L50, further underscoring the complexity of pre-miR21 regulatory mechanisms and its impact on “mature miR” formation. Through its ability to suppress the network of interaction within the RNA strand, we show how this peptide can modulate pre-miR21’s accessibility to the downstream binding partner (i.e., the Dicer protein). At the same time, L50 displayed a limited influence on pre-miR21’s FES, unable to induce any significant shift in the “stacked-in/bulged-out” equilibrium. This result not only helps us rationalize the weak inhibitory power of a nanomolar binder like L50, but also points to a clear strategy for improving its efficacy. While our strategy helped us navigate the FES of a specific microRNA, there is still room for improvement in developing a generalized platform for designing potent RNA binders. Indeed, it is worth stressing that during our OneOPES simulations of the holo-complex, we assumed that L50 retains the same binding mode in both the “bulged-out” and “stacked-in” states (as suggested by the NMR structures resolved in PDB ID 5UZZ). If this assumption proves incorrect, additional CVs would need to be introduced to capture the ligand’s dynamics and handle variations in the binding mode more effectively. Additionally, to account for the effects of different ionic strengths, we carried out an unbiased MD simulation of the L50/pre-miR21 complex under physiological NaCl concentration. While in the present case no substantial differences compared to our prior holo-pre-miR21’s simulation were observed (see Fig. S7), future studies should consider this aspect when designing more accurate simulations.

Concluding, peptides are making a comeback as effective therapeutic candidates for various diseases^41 42 43^. Our findings and the structure we have released provide valuable insights for designing new therapeutic peptides (or small molecules) targeting the miR21 regulatory pathways, with the auspice of identifying novel lead-compounds able to both suppress RNA’s motility and strongly stabilise the “bulged-out” state. These new therapeutics could have potential applications in disease intervention and precision medicine. Moreover, since our strategy is not tailored to the specific system under study, it can be easily adapted to a variety of miR-ligand complexes, overcoming the typical difficulties encountered by other in-silico methods in tackling the high mobility of miRs and there is a need for better and newer techniques in this field.

Moving forward, continued investigation into the molecular mechanisms underlying miR dynamics will be essential for harnessing their therapeutic potential and unraveling the complexities of gene regulation in health and disease.

## Data Availability

The PDB structure of the L50/pre-miR21 heterocomplex in the “bulged-out” state is released as part of the Supplementary Material. The input files to replicate all the simulations can be found on https://github.com/valeriorizzi/miRNA and on PLUMED NEST^44^.

## Associated content

Supporting Information is available, reporting additional computational details on both the classical MD simulations and the enhanced sampling trajectories. The structure of the L50/pre-miR21 complex in the “bulged-out” state is released as a PDB in the supplementary material.

## Acknowledgements

The work was financially supported by the Swiss National Science Foundation and Bridge funding schemes (project numbers: 200021 204795, CRSII5 216587, and 40B2-0 203628). We acknowledge PRACE and the Swiss National Supercomputing Centre for supercomputer time allocations on Piz Daint (project ID: s1228). The authors are grateful to Nicola Piasentin for designing the table of contents figure.

## Notes

### Competing Interest Statement

The authors have declared no competing interest.

### Summary of Updates

-Additional analyses on the apical loop of pre-miR21 -Analysis on L50's conformational plasticity -PCA on apo and holo MD simulations

