## Supplementary Information for "Investigating Ligand-Mediated Conformational Dynamics of Pre-miR21: A Machine-Learning-Aided Enhanced Sampling Study"

### SUPPLEMENTARY MATERIAL

The data hereby reported are divided into the following sections:

- Supplementary Data 1: Additional details on the apo- and holo-pre-miR21 MD simulations;
- Supplementary Data 2: Cluster analyses on the apo- and holo-pre-miR21 MD simulations;
- Supplementary Data 3: PCA on apo- and holo-pre-miR21 MD simulations;
- Supplementary Data 4: Details on apo-pre-miR21 OneOPES simulation;
- Supplementary Data 5: Details on holo-pre-miR21 OneOPES simulation;
- Supplementary Data 6: Binding mode of L50 in the "bulged-out" state;
- Supplementary Data 7: Comparison of "bulged-out" conformations between the apo- and holo-pre-miR21 OneOPES simulations;
- Supplementary Data 8: Details on holo-pre-miR21 MD simulation at salinity 0.15 NaCl.

### Supplementary Data 1

In the following section, we present additional data supporting our analysis on both apo-pre-miR21 and holo-pre-miR21 MD simulations. We carried out an RMSD analysis of the apical loop in both systems, highlighting its flexibility (see Fig. S1a and S1b). Additionally, Fig. S1c presents the RMSD of the cyclic peptide L50, calculated based on its Cαs, while Fig. S1d shows the time evolution of intra-peptide hydrogen bond distances in L50. By comparing the fluctuations in L50's RMSD with its intra-peptide H-bond distances, we can speculate that these fluctuations are primarily due to internal motions of the peptide, where certain hydrogen bonds are broken and re-formed over time.

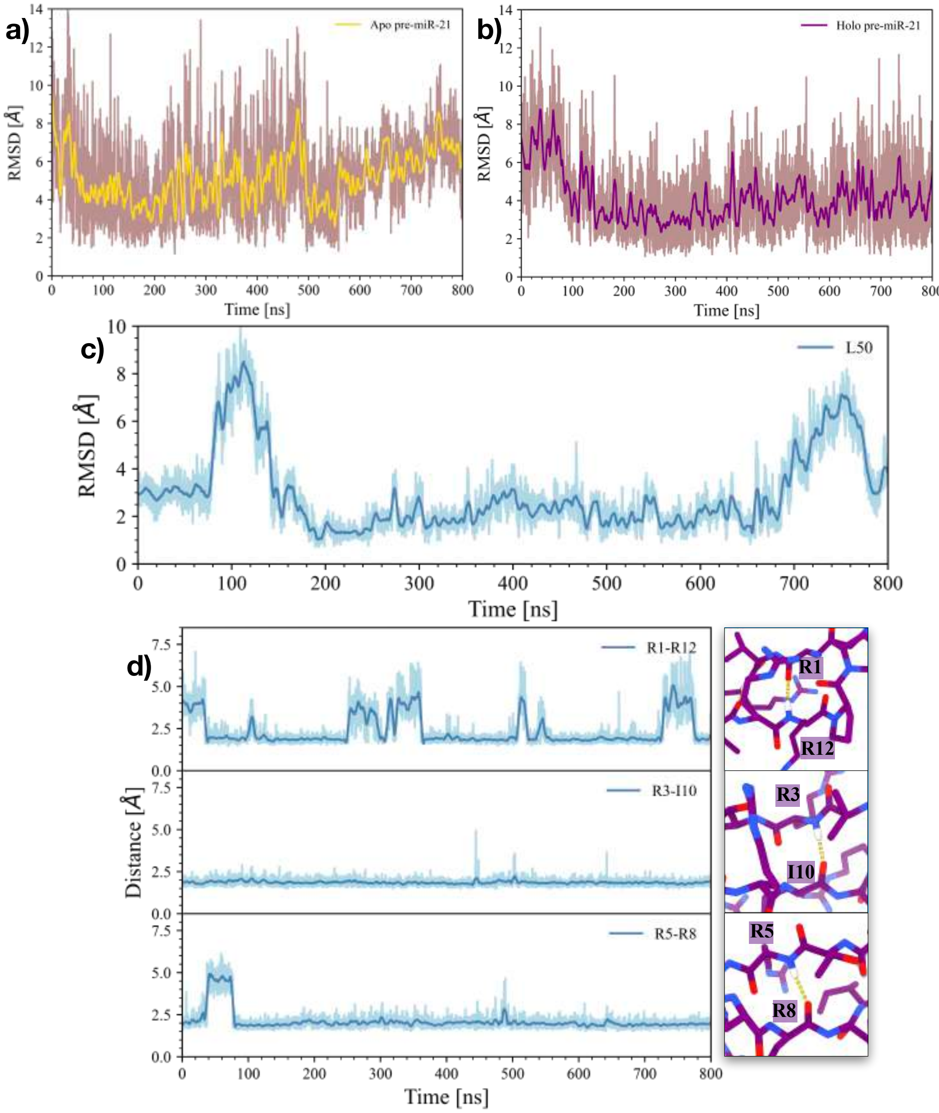


Figure S1: **RMSD analysis of pre-miR21 apical loop and L50 cyclic peptide dynamics. a-b)** RMSD of the apical loop in the apo- (a) and holo-pre-miR21 (b) MD simulations. The running averages are colored in yellow and purple, respectively. **c)** RMSD of the cyclic peptide L50, calculated based on the $\alpha$ carbons. **d)** Time evolution of the intra-peptide hydrogen bond distances in L50 during the simulations. The residues involved in the H-bonds are depicted on the panels on the right side.

#### Supplementary Data 2

In the following section, we present the supplementary data generated to support our investigation of the cluster analyses we performed on both apo-pre-miR21 and holo-pre-miR21 MD simulations. The results, as shown in Fig. S2a, highlight the flexibility of the apo-pre-miR21 system, which allows us to observe a transition event involving the conformational change of A29. This transition mainly results in two distinct cluster families, i.e. one for the "stacked-in" state and a second one for the "bulged-out" state. In contrast, the limited plasticity of the holo-pre-miR21 system prevents us from observing any conformational change of A29 within the 800 ns MD simulation (see Fig. S2b).

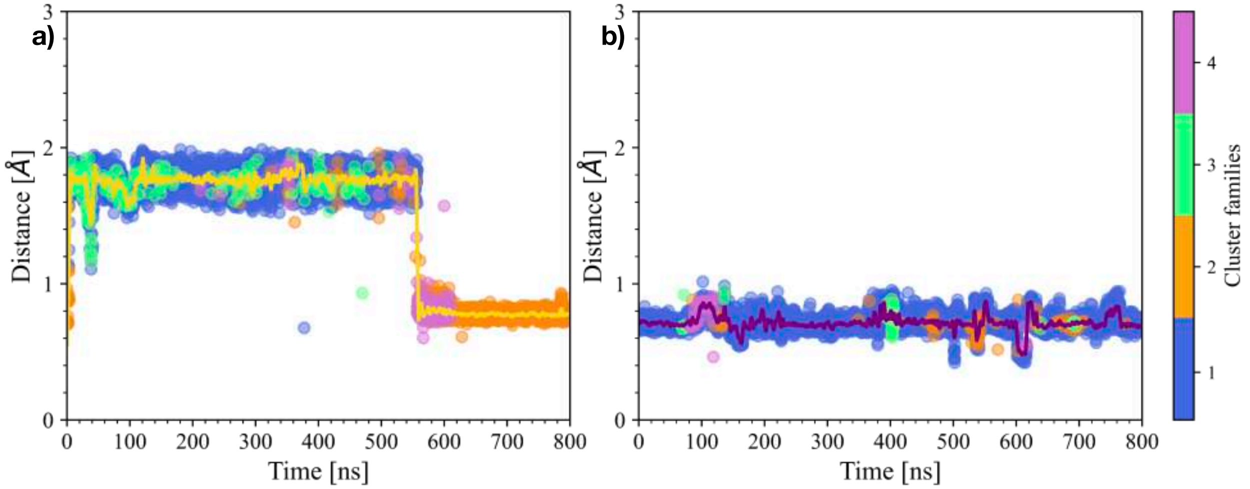


Figure S2: **Additional details about the cluster analyses carried out on the apo- and holo-pre-miR21 MD simulations. a-b)** Time evolution of the distance between A29 and the nucleobases on the opposite RNA strand (i.e., G45 and C46) for the apo- (a) and holo-pre-miR21 (b) MD simulations. Each point is colored according to its corresponding cluster family, as indicated by the color bar on the right hand side and illustrating the flexibility of the apo-pre-miR21 system with respect to the holo-pre-miR21. The running averages are colored in yellow and purple, respectively.

#### Supplementary Data 3

In this section, we present the results of the principal component analysis (PCA) performed on both apo-pre-miR21 and holo-pre-miR21 MD simulations. Porcupine plots corresponding to the first eigenvector were generated for each system, revealing the primary modes of motion. In detail, Fig. S3a and Fig. S3b show the direction of the first eigenvector, highlighting that the presence of L50 suppresses the anti-correlated motion between the regions *G28-U31* (yellow) and *C39-A50* (green). Moreover, the Scree plots on the right hand side display the percentage of variance explained by the eigenvectors, highlighting the contributions of each mode to the overall conformational dynamics of the apo and holo systems.

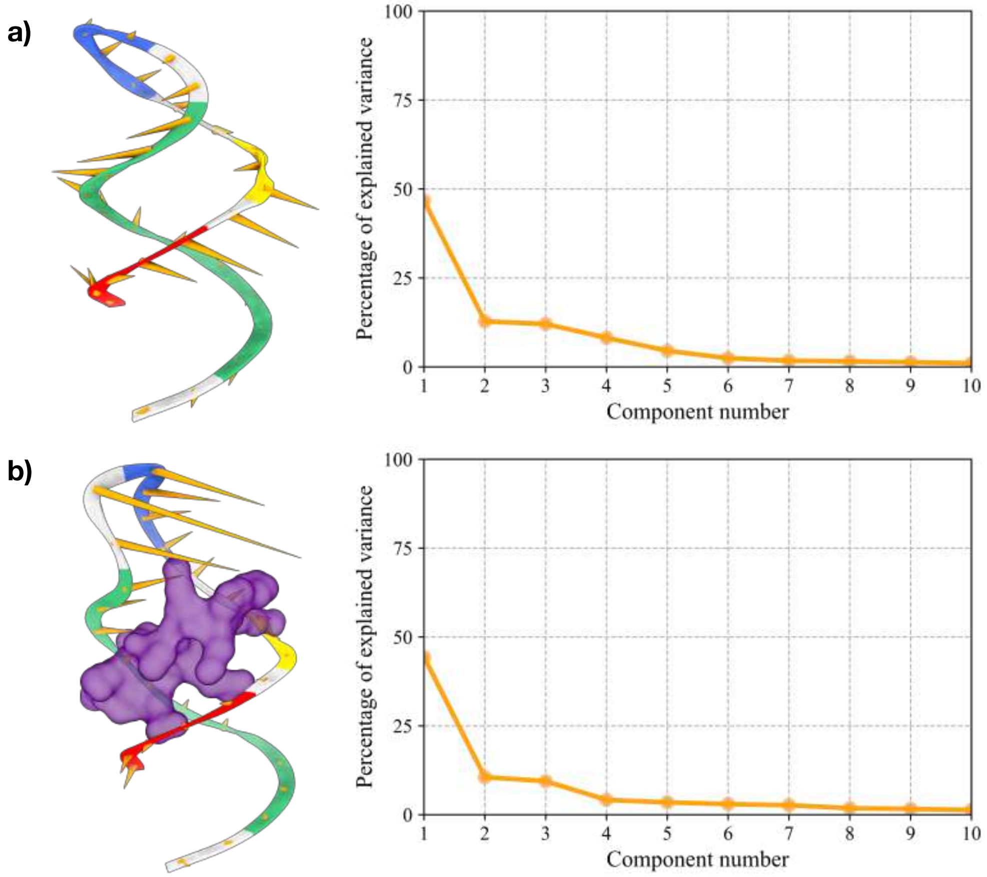


Figure S3: **Principal component analysis carried out on the apo- and holo-pre-miR21 MD simulations. a-b)** Porcupine plots of the first eigenvector, as calculated for both the apo- (a) and holo-pre-miR21 (b) MD simulations. Each macroregion of pre-miR21 is colored according to the definition in Fig. 3. On the right side, scree plots displaying the percentage of explained variance carried out by the eigenvectors for both systems.

#### Supplementary Data 4

In the following section, we report all the supplementary data we generated to support our investigation on the apo-pre-miR21's OneOPES simulation.

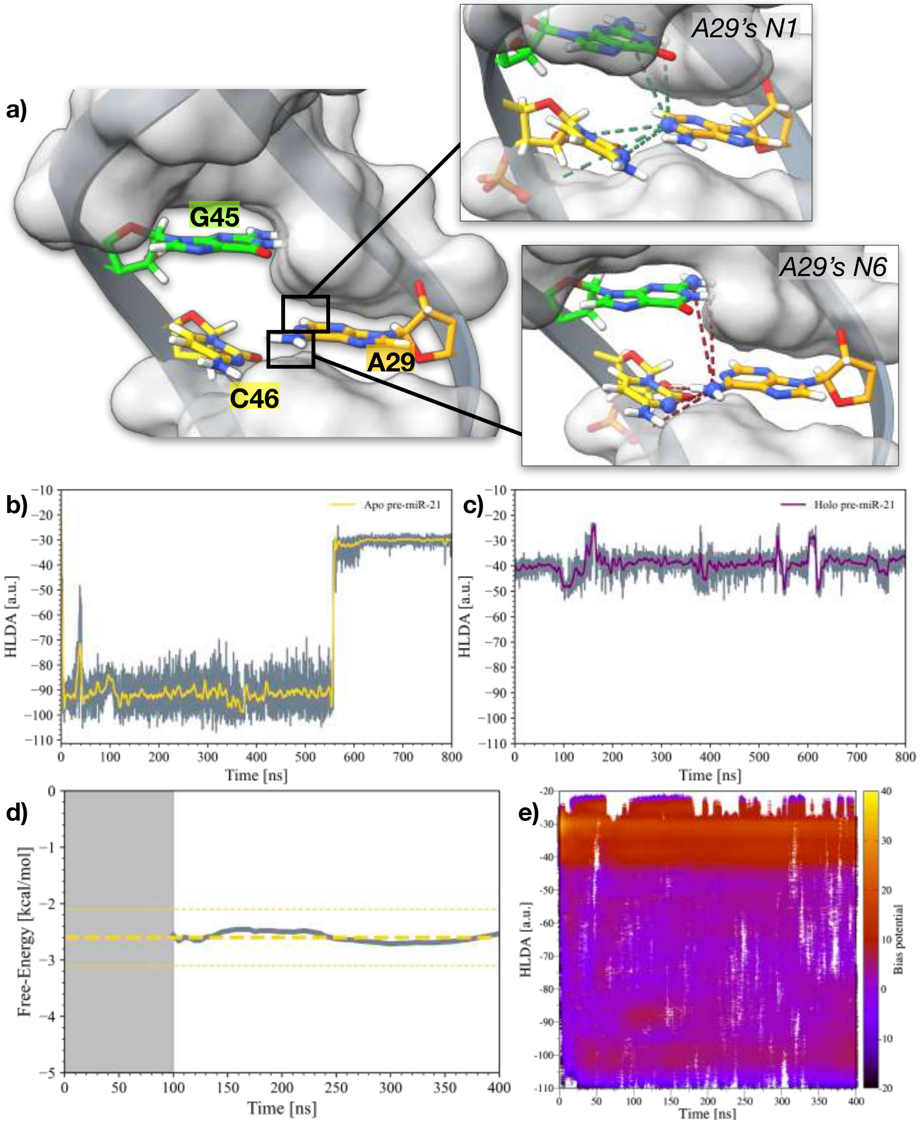


Figure S4: **Additional details about the apo-pre-miR21's OneOPES simulation. a)** The three nucleobases among which we built the HLDA CV. The contacts established by A29's N1 and N6 are reported in the top and bottom insets, respectively. A29 is colored in orange, G45 in green, and C46 in yellow. The phosphate backbone of the nucleotides is colored in grey. **b)** Reweighted HLDA CV for the apo-pre-miR21 system, based on unbiased MD simulation. **c)** Reweighted HLDA CV for the holo-pre-miR21 system, based on unbiased MD simulation. **d)** ΔG of A29's conformational change as a function of the simulation time for apo-pre-miR21's OneOPES simulation. The ΔG over time is colored in grey, whereas the average ΔG value is represented through a yellow dashed line. **e)** Plot displaying the exploration of the whole HLDA CV during the OneOPES simulation. Each point is represented by a colour that corresponds to the associated bias potential, as indicated by the colour bar on the right side.

#### Supplementary Data 5

The following section reports all the supplementary data we generated during the holo-pre-miR21's OneOPES simulations.

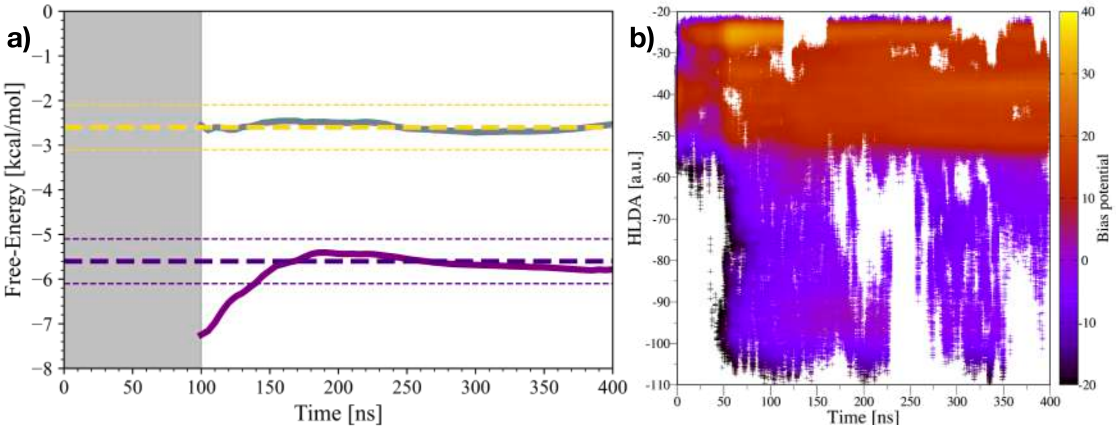

Figure S5: **Additional details on the OneOPES simulations carried out on the holo-pre-miR21 hetero-complex. a)** ΔG of A29's conformational change as a function of the simulation time for apo-pre-miR21's (grey) and holo-pre-miR21's (purple) OneOPES simulations. For the former, the average ΔG value is represented through a yellow dashed line, while it is displayed as a purple dashed line for the latter. **b)** Plot displaying the exploration of the whole HLDA CV during holo-pre-miR21's OneOPES simulation. Each point is represented by a colour that corresponds to the associated bias potential, as indicated by the colour bar on the right side.

#### Supplementary Data 6

In this section, we explore the molecular interactions established by the L50/pre-miR21 heterocomplex in the "bulged-out" state, focusing on the specific binding interactions illustrated in Fig. 5 and their respective distances. Together with the estimate of the ΔG, the distances measured between interacting residues offer a quantitative understanding of how the cyclic peptide L50 associates with pre-miR21, thereby leading the way about how to improve L50's inhibitory power.

| *L50* | *pre-miR21* | *Distance [Å]* |
|---|---|---|
| Arg1 | G28 | 1.9 |
| Arg1 | A29 | 2.9 |
| Arg3 | U26 | 1.9 |
| Thr4 | G45 | 1.9 |
| Arg5 | G25 | 2.0 |
| Lys7 | G44 | 1.7 |
| Lys7 | G45 | 1.8 |
| Arg8 | G23 | 1.8 |
| Arg8 | U24 | 1.8 |
| Arg9 | G44 | 2.0 |
| Arg11 | U40 | 1.8 |
| Arg11 | C41 | 2.0 |
| Arg12 | G25 | 1.9 |
| Arg12 | U26 | 2.0 |

Table S1: Table depicting the L50/pre-miR21 interactions portrayed in Fig. 5, where we showed the representative structure of holo-pre-miR21's "bulged-out" state.

#### Supplementary Data 7

In this section, we present a detailed comparison between the "bulged-out" conformational states of pre-miR21, as observed in the OneOPES simulations of both the apo- and holo-pre-miR21 systems. By analyzing the structural differences between these two states, we aim to highlight how the presence of the L50 peptide influences the conformational dynamics of A29 and the surrounding RNA regions.

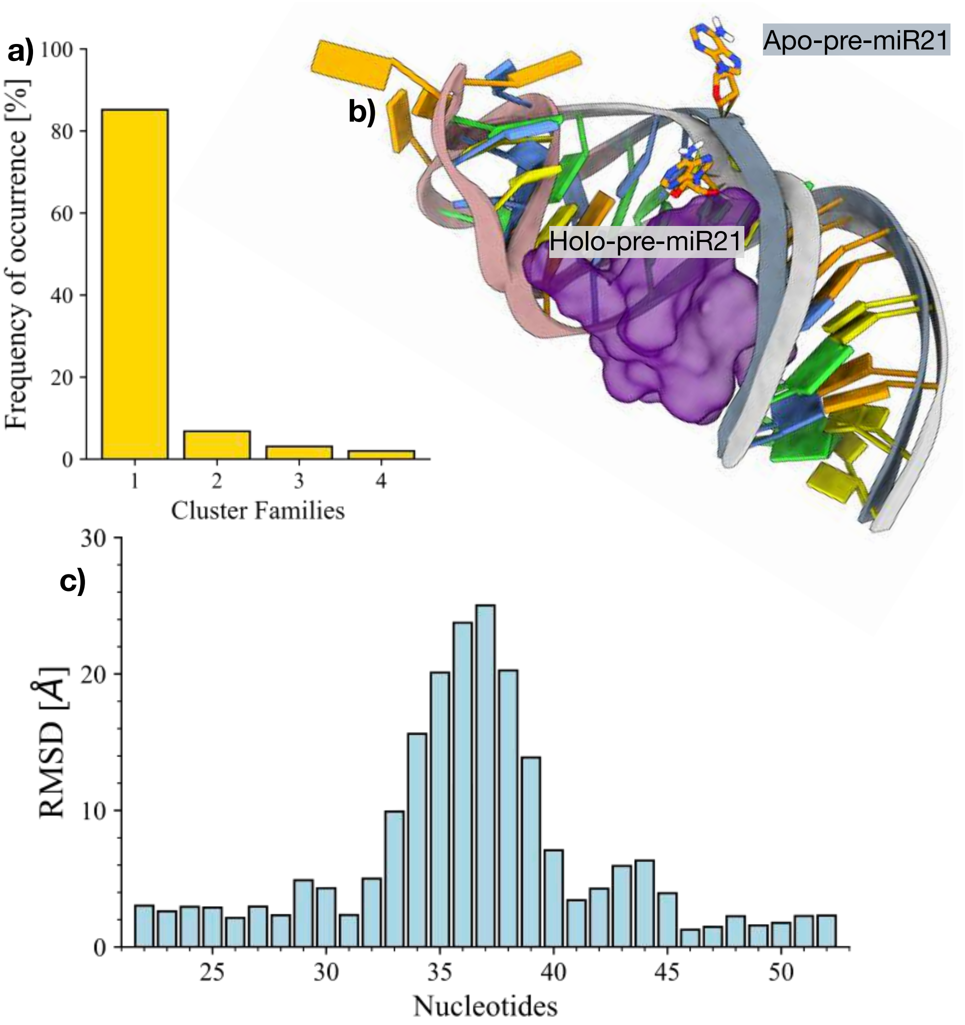


Figure S6: **Comparison of the bulged-out states of pre-miR21 from the apo- and holo- OneOPES simulations. a)** Cluster analysis performed on the extrapolated frames from the bulged-out basin in the apo-pre-miR21 systems. **b)** Structural superimposition of the bulged-out state obtained from the apo-pre-miR21 and holo-pre-miR21 OneOPES simulations, demonstrating the differences in RNA conformation influenced by the presence of the L50 peptide. **c)** Per-residue RMSD plot comparing the bulged-out conformations from the apo and holo OneOPES simulations, showing the structural variability at specific nucleotide positions.

### Supplementary Data 8

The following section presents the supplementary data generated from the unbiased MD simulation of the holo-pre-miR21 system, carried out in the presence of a physiological concentration of NaCl (0.15 M). This simulation was performed to explore how the ionic environment influences the dynamics and stability of the RNA-peptide complex. The results provide insight into the binding interactions and conformational stability under conditions closely mimicking physiological ionic strength.

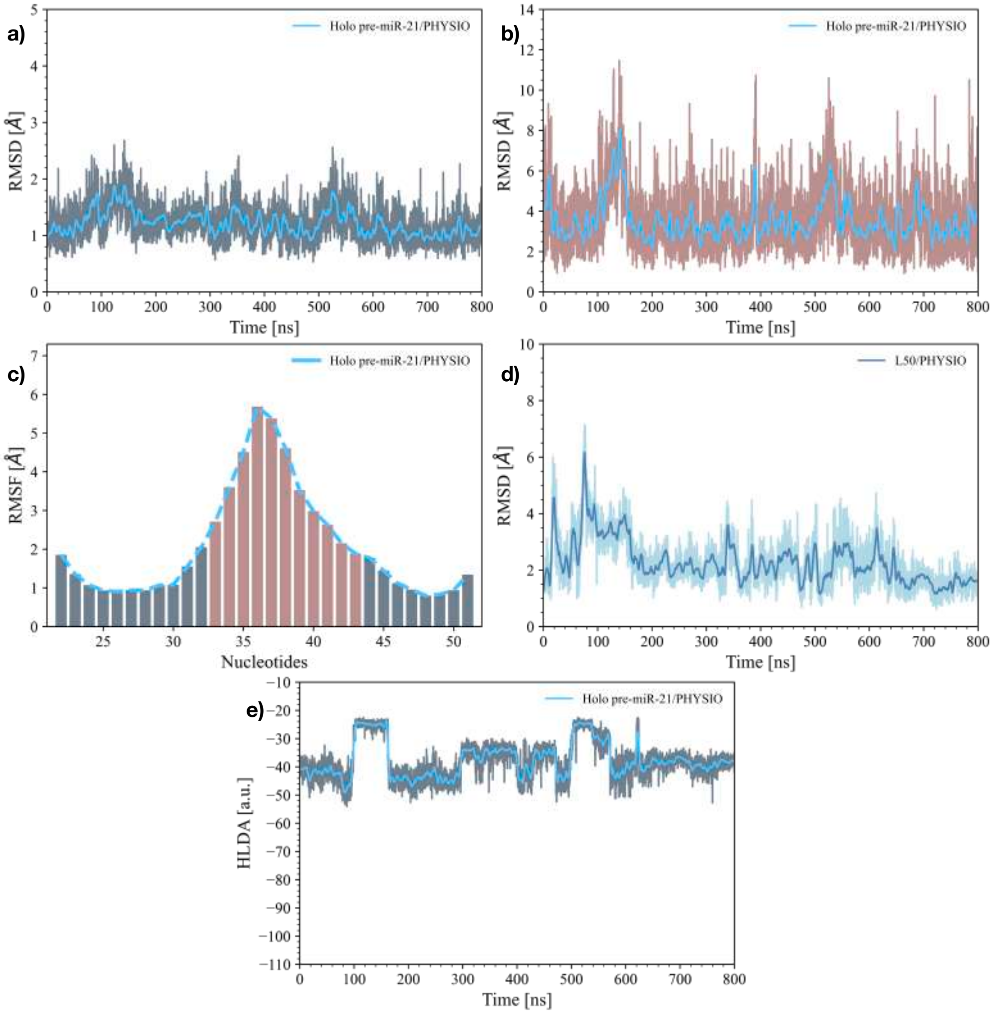


Figure S7: **Details of the MD simulations on the holo-pre-miR21 heterocomplex at physiological salinity level. a)** RMSD of pre-miR21's backbone in the holo- MD simulation. **b)** RMSD of the apical loop in the holo-pre-miR21 MD simulations. In both (a) and (b), the RMSD running averages are superimposed in azure. **c)** Histograms displaying the per-nucleobase RMSF for the holo-pre-miR21 system. **d)** RMSD of the cyclic peptide L50, calculated based on the $\alpha$ carbons. **e)** Reweight of the HLDA CV on the holo-pre-miR21's unbiased MD simulation.
